# Altered markers of brain metabolism and excitability are associated with executive functioning in young children exposed to alcohol *in utero*

**DOI:** 10.1101/2024.05.06.592786

**Authors:** Meaghan V. Perdue, Mohammad Ghasoub, Madison Long, Marilena M. DeMayo, Tiffany K. Bell, Carly A. McMorris, Deborah Dewey, W. Ben Gibbard, Christina Tortorelli, Ashley D. Harris, Catherine Lebel

## Abstract

Prenatal alcohol exposure (PAE) is the leading known cause of birth defects and cognitive disabilities, with impacts on brain development and executive functioning. Abnormalities in structural and functional brain features are well-documented in children with PAE, but the effects of PAE on brain metabolism in children have received less attention. Levels of brain metabolites can be measured non-invasively using magnetic resonance spectroscopy (MRS). Here, we present the first study of PAE-related brain metabolite differences in early childhood (ages 3-8 years) and their associations with cognitive performance, including executive functioning (EF) and pre-reading skills. We measured metabolites in two cohorts of children with PAE and unexposed children using MRS in the anterior cingulate cortex (ACC; cohort 1) and left temporo-parietal cortex (LTP; cohort 2). Total choline (tCho), a marker of membrane/myelin metabolism, was elevated in both regions in children with PAE compared to unexposed children, and glutamate+glutamine (Glx), a marker of excitability, was elevated in the ACC. The PAE group exhibited more difficulties with EF, and higher tCho was associated with better EF in both PAE and unexposed groups. In addition, elevated Glx in the ACC was associated with poorer inhibitory control within the PAE group only. LTP metabolites were not significantly associated with pre-reading skills in PAE or unexposed groups. Together, these findings point to altered membrane metabolism and excitability in young children with PAE. These findings provide new insight to potential mechanisms by which PAE disrupts brain development and cognitive functioning in early childhood.

## Introduction

Alcohol consumption during pregnancy can lead to a range of cognitive, behavioral, and neurological difficulties in offspring (Popova et al., 2023). Prenatal alcohol exposure (PAE) is the leading known cause of developmental disabilities, with approximately 10% of children exposed worldwide (Popova et al., 2023). Alcohol is a teratogen that can easily cross the placenta, disrupting the fetus’ neurodevelopment at any stage. Disruptions include, but are not limited to, altered neuronal proliferation and migration (Almeida et al., 2020; Veazey et al., 2013), synaptogenesis (Adams et al., 2023; Basavarajappa & Subbanna, 2023), dendritic spine morphology (Cui et al., 2010; Lee et al., 2021), and gliogenesis and myelination (Cantacorps et al., 2017; Darbinian & Selzer, 2022), including phospholipid metabolism (Darbinian et al., 2023; Hwang et al., 2023).

The teratogenic effects of alcohol on brain development can lead to long-term cognitive and behavioral difficulties (Mattson et al., 2019; Subramoney et al., 2018). Children with PAE may be diagnosed with Fetal Alcohol Spectrum Disorder (FASD), which is characterized by physical and neurodevelopmental impairments, though diagnostic criteria vary (Cook et al., 2016; Popova et al., 2023). Children with PAE often exhibit attention-deficit/hyperactivity disorder (ADHD)-like behavior (Pinner et al., 2023), along with executive and social functioning challenges (Khoury et al., 2015; Kodituwakku et al., 2001; Mattson et al., 2019; Mattson & Riley, 1999; Rasmussen et al., 2007, 2013; Rasmussen & Bisanz, 2009; Rockhold et al., 2021). In addition, children with PAE are more likely than their unexposed peers to have difficulties in academic performance, including reading and math (Kodituwakku et al., 2011; O’Leary et al., 2013). Academic difficulties experienced by children with PAE may arise from domain-general cognitive difficulties, including problems in memory, attention, processing speed and executive functions (Kodituwakku et al., 2011).

These behavioral difficulties emerge in the context of PAE-related brain differences (Donald et al., 2015; Lebel et al., 2011; Moore & Xia, 2022). Reduced white matter and gray matter volume (Nardelli et al., 2011; Subramoney et al., 2022), altered microstructure and developmental trajectories (Donald et al., 2024; Gimbel et al., 2023; Kar et al., 2021, 2022; M. Long et al., 2024; X. Long & Lebel, 2022), and disrupted structural and functional connectivity (X. Long et al., 2019, 2020; Roos et al., 2021; Wozniak et al., 2013) have been reported in children and adolescents with PAE. Moreover, neural alterations are observed even with relatively low levels of PAE, around 1 drink/week (X. Long & Lebel, 2022).

Despite known impacts of PAE on brain development, the neurobiological mechanisms underlying cognitive difficulties in children with PAE remain poorly understood. Examining features of brain metabolism and chemistry provides complementary evidence to structural and functional neuroimaging data, offering a window to biochemical mechanisms underlying altered brain development, metabolism, and functioning in those with PAE. Brain metabolites are naturally occurring chemicals that are crucial for development, metabolism, and neural signalling that can be measured non-invasively using magnetic resonance spectroscopy (MRS). Five key brain metabolites of interest that are measured using standard MRS methods include total NAA (N-acetylaspartate + N-acetylaspartylglutamate; tNAA), which reflects neuronal health and metabolism, total creatine (creatine + phosphocreatine; tCr), which reflects energy homeostasis, total choline (phosphocholine + glycerophosphocholine + choline, tCho), which reflects phospholipid metabolism; glutamate + glutamine (Glx), which reflects neural excitability, and myo-inositol (Ins) a marker of glial tissue. For an in-depth discussion of the roles of each metabolite in the brain, see Rae (2014).

MRS offers insight to potential neurochemical and metabolic mechanisms by which alcohol disrupts brain structure development and functioning. Specifically, it has been posited that alcohol disrupts the cortico-striatal excitation/inhibition balance by increasing glutamatergic activity and supressing migration of inhibitory neurons to their cortical targets (Bariselli & Lovinger, 2021). This system can be investigated in humans using MRS to measure Glx and g-aminobutyric acid (GABA). In addition, myelination is impaired in PAE, and preclinical models point to disruption of oligodendrocyte proliferation, migration, and maturation (Darbinian & Selzer, 2022). However, the mechanism by which alcohol impacts myelination is not fully understood. The tCho signal in MRS is a marker of phospholipid metabolism, including the synthesis/repair and break-down of neuronal cell membranes and myelin (Rae, 2014), which may be impacted by PAE.

Despite the potential of MRS to provide insight to neurometabolic brain alterations, only six studies have applied MRS to investigate PAE (Astley et al., 2009; Cortese et al., 2006; du Plessis et al., 2014; Fagerlund et al., 2006; Gonçalves et al., 2009; Howells et al., 2016), and half of these included sample sizes of 10 or fewer in each group. Nonetheless, several trends have emerged in this small body of literature, indicating altered brain metabolism and excitability in infants and youth with PAE relative to controls. Reduced tCho in children and adolescents with PAE has been reported in three studies that measured metabolites in different regions, with effects shown in frontal parietal white matter, left striatum, and cerebellum, indicating that tCho may be affected globally in PAE (Astley et al., 2009; du Plessis et al., 2014; Gonçalves et al., 2009). Lower tCho in PAE is thought to indicate altered phospholipid metabolism, impaired myelin maintenance, and reduced cell density/arborization relative to unexposed groups. One study linked higher rates of alcohol exposure during pregnancy with lower tCho and tNAA, and higher Glx, in children ages 8-17 years, highlighting the persistent effects of PAE on neurometabolic outcomes into adolescence (du Plessis et al., 2014). Glx differences in PAE are mixed across studies: du Plessis and colleagues (2014) found that elevated cerebellar Glx levels in children/adolescents were associated with increased rates of alcohol consumption around conception and during pregnancy, while Howells and colleagues (2016) found that parietal white matter Glx levels were reduced in newborns with PAE relative to controls. These contrasting patterns may reflect differences across studies in developmental stages examined, brain regions and tissue types from which metabolites were measured, and relative contributions of glutamate and glutamine to the Glx signal. These varied methods and findings highlight the need for further investigation across different age groups and brain regions, to better characterize the impacts of PAE on neurochemistry across development. Moreover, no study to date has investigated associations between metabolites and cognitive traits in participants with PAE.

Here, we present the first study of neurometabolic differences associated with PAE in early childhood, and report novel links between brain metabolites and cognition in this population. We measured brain metabolites and cognitive traits in two cohorts of young children (ages 3-8 years) with PAE in comparison to unexposed controls. Metabolite levels (tCho, tNAA, tCr, Glx, & mI) were measured in the anterior cingulate cortex (ACC; Cohort 1) and left temporo-parietal cortex (LTP; Cohort 2). These regions were selected based on their roles in the EF (ACC) and language/pre-reading (LTP) skills targeted in the study; due to time constraints, we were not able to collect MRS data from multiple voxels within study visits. Children’s EF (Cohort 1) and pre-reading skills (Cohort 2) were assessed using age-appropriate standardized measures. We predicted that children with PAE would have lower tCho and higher Glx relative to controls, in line with prior research (Astley et al., 2009; du Plessis et al., 2014; Gonçalves et al., 2009); our analyses of tNAA, tCr and myo-inositol (mI) were exploratory given the lack of prior research in this population. In addition, we predicted that tCho and Glx in the ACC would be associated with EF, while tCho and Glx levels in the LTP would be associated with pre-reading skills.

## Methods

### Participants

This study included children with PAE from a longitudinal study of early childhood brain development in children with PAE, and unexposed children drawn from a longitudinal study of typical brain development with matching brain imaging protocols. Children with PAE were recruited from Alberta Children’s Services, caregiver support groups, and early intervention services. PAE was confirmed from the children’s welfare file and/or from caregivers, caseworkers, and family members. All children were in adoptive, foster, or kinship care. None of the children had a formal FASD diagnosis at the time of assessment given their young age, as FASD diagnoses in some parts of Alberta typically begins at 7 years of age. All children with PAE were born at ≥33 weeks’ gestation and spoke English as their primary language. Children with epilepsy, autism spectrum disorder, history of head injury, MRI contraindications, and any other genetic or medical conditions associated with cognitive disabilities were excluded from the study. Given their high co-occurrence with PAE, children with conditions such as ADHD, learning disabilities, and other mental health diagnoses were included, numbers of participants with these conditions included in the current study are listed in Table S1. Further details on the children in this study are reported in (Kar et al., 2021). Children without alcohol exposure (henceforth, “unexposed children”) were recruited from the community in Calgary, Alberta as well as the Alberta Pregnancy Outcomes and Nutrition (APrON) study. All unexposed children were born at ≥35 weeks gestation and spoke English as their primary language. Inclusion criteria included no MRI contraindications, history of brain injury, or genetic, motor, or developmental disorders at the time of enrollment. Several participants received diagnoses of developmental disorders (ADHD, speech/language difficulties, learning disabilities) over the course of longitudinal study participation; numbers are listed in Table S1. These children had no PAE as verified by prospective maternal report. Further details on this study are reported in (Reynolds et al., 2020). The sample of unexposed children in this study were included in a larger study of metabolite development across childhood (Perdue et al., 2023); the present study includes a subset of these children within age ranges restricted to match the age ranges of the participants with PAE for each cohort. Parent/guardian written informed consent and child verbal assent were obtained for each child. Both studies were approved by the University of Calgary’s Conjoint Health Research Ethics Board (REB14-2266, REB13-0020).

Both studies are accelerated longitudinal studies of brain development, and MRS data was acquired from the ACC during the first wave of data collection, and from the LTP in subsequent longitudinal study visits. Children were therefore divided into two cohorts for the present study based on the availability of MRS data acquired from the ACC (Cohort 1) and the LTP (Cohort 2). Longitudinal data acquired within each region was included from some individuals as available; longitudinal data points were acquired a minimum of 6 months apart. There is some overlap in the participants included in each cohort, but MRS data were acquired from the ACC and LTP at different study visits for all participants. All available good quality MRS data was included in analysis to maximize the stability of our estimates and the statistical power of our models.

Cohort 1 (ACC) consisted of 21 datasets collected from 20 children (11 females) with PAE (5.38 ± 1.11 years) and 76 datasets collected from 72 unexposed children (33 females; 4.51 ± 0.950 years). Cohort 2 (LTP) consisted of 28 datasets collected from 24 subjects (16 females) with PAE (6.45 ± 0.764 years) and 128 datasets from 71 unexposed children (34 females; 6.45 ± 0.925 years). These numbers reflect the participants/datasets included in analysis after exclusion for data quality issues (detailed below). Ten children with PAE and 39 unexposed children had MRS data from both regions and were included in both cohorts, though MRS data were acquired from each region at different study visits (6 months - 1 year apart).

### Metabolite data acquisition

Brain metabolites were measured using magnetic resonance spectroscopy (MRS) included in a multimodal magnetic resonance imaging (MRI) protocol. MRI and MRS data were acquired with a 3T GE MR750w MR system with a 32-channel head coil at the Alberta Children’s Hospital (Calgary, AB, Canada) by staff who are highly skilled in pediatric neuroimaging. Children were scanned while watching a movie or during natural sleep; no sedation was used. T1-weighted anatomical images were acquired using a spoiled gradient echo sequence (210 axial slices; 0.9 x 0.9 x 0.9mm resolution, TR = 8.23 ms, TE = 3.76 ms, flip angle = 12°, matrix size = 512 x 512, inversion time = 540 ms). These images were reconstructed to provide axial, sagittal, and coronal views at the scanner which were used for placement of spectroscopy voxels. MRS data were acquired using short echo time Point RESolved spectroscopy (PRESS; TE = 30 ms, TR = 2000 ms, 96 averages, 20 x 20 x 15 mm^3^ voxel size). A strong body of literature demonstrates the test-retest reliability and stability of PRESS sequences and supports the validity of MRS measurements for longitudinal research (Baeshen et al., 2020; Fayed et al., 2009; Gasparovic et al., 2011; Soreni et al., 2010; Volk et al., 2018, 2019). MRS voxels were placed in either the ACC or in the LTP by trained members of the research team according to detailed instructions and reference images. The ACC voxel was localized anterior to and at approximately the same level as the genu of the corpus callosum viewed on a midsagittal slice and consisted almost entirely of gray matter (Figure 1). The LTP voxel was localized to capture the left angular gyrus based on all three image planes, and included portions of the supramarginal gyrus, parietal operculum, and posterior superior temporal gyrus due to the extent of the voxel (Figure 1). The LTP voxel consisted of gray and white matter, excluding CSF to the extent possible.

**Figure 1.**
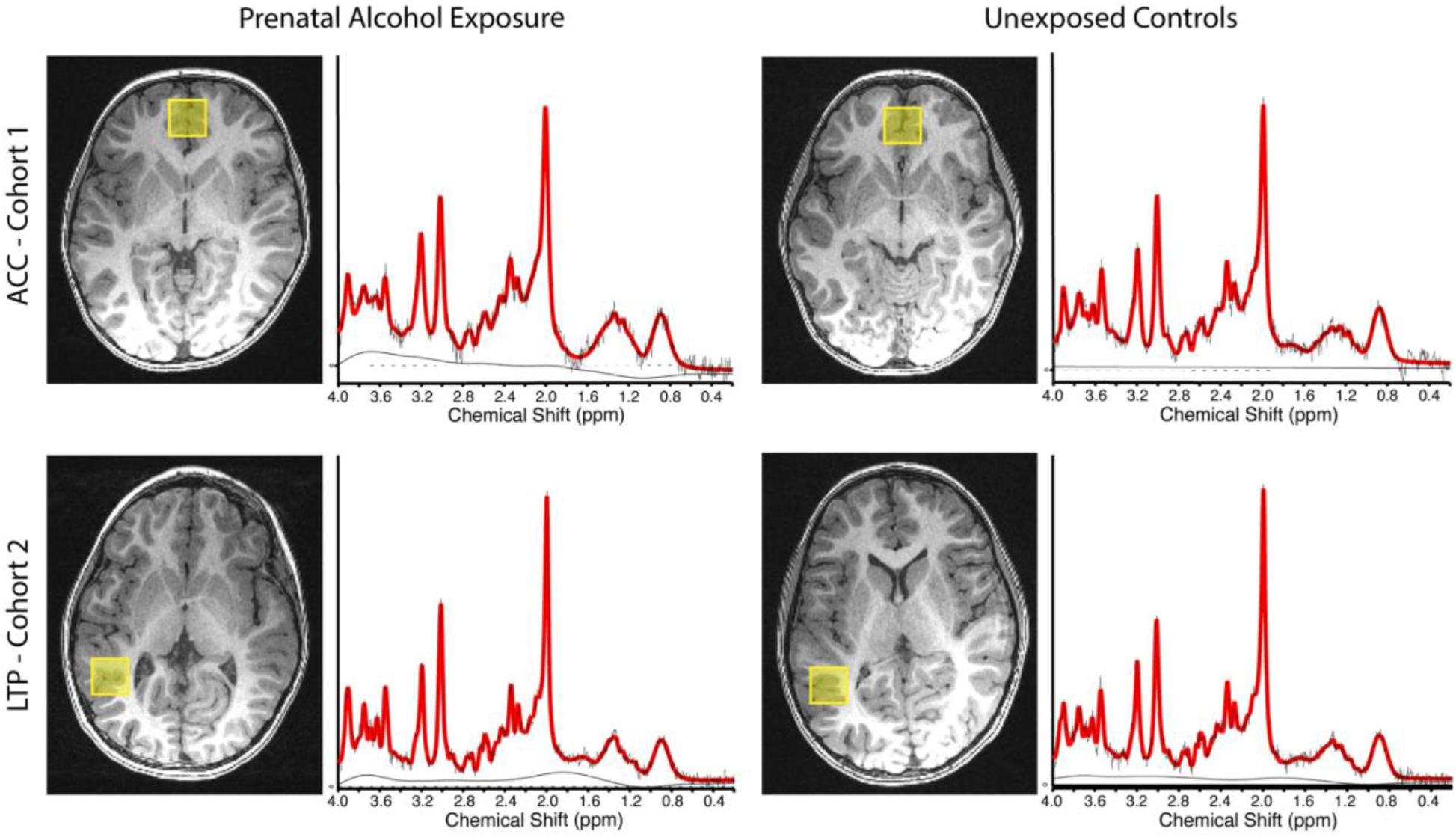
MRS voxel positions and sample spectra. ACC shown on top row, LTP shown on bottom row, participants with PAE shown on left, unexposed participants shown on right. Sample participant age and sex are as follows: Cohort 1 PAE: 4.87 year-old male, Cohort 1 unexposed: 4.89 year-old male, Cohort 2 PAE: 6.14 year-old female, Cohort 2 unexposed: 6.25 year-old female

### Metabolite data processing and quantification

The PRESS acquisition was pre-processed with the FID-A toolbox (Simpson et al., 2017). Pre-processing included coil combination, removal of bad averages, frequency alignment, and phase correction, according to recent recommendations (Near et al., 2020). Metabolites were then fit using LCModel v6.3 (Provencher, 1993). The metabolites of interest were: tNAA, tCr, tCho, mI and Glx. The basis set used for quantification included alanine, ascorbate, aspartate, choline, citrate, creatine, ethanol, GABA, glycerophosphocholine, glutathione, glucose, glycine, glutamine, glutamate, water, mI, lactate, NAA, NAAG, phosphocholine, phosphocreatine, phosphoryl ethanolamine, scyllo-inositol, taurine and beta-Hydroxybutyrate, simulated using the FID-A toolbox (Simpson et al., 2017) with sequence specific timings and RF pulse shapes. Fitted spectra were visually inspected for quality assurance and datasets with poor spectra were excluded from analysis (ACC [Cohort 1]: PAE n=5, unexposed n=3; LTP [Cohort 2]: PAE n=0, unexposed n=3). Quantitative quality metrics (signal-to-noise ratio [SNR] and linewidth [LW] for NAA) were obtained using the “op_getLW.m” and “op_getSNR.m” functions in FID-A. The Gannet CoRegStandAlone function (Harris et al., 2015), which calls SPM12 segmentation (Ashburner & Friston, 2005), was used to quantify tissue content within the MRS voxels by co-registering them to the T1-weighted image acquired during the same scanning session. Co-registration and segmentation outputs were visually inspected to ensure accurate localization and segmentation of the MRS voxels. Absolute metabolite values are expressed in molal units (moles/kg), as per the approach recommended in the MRS consensus statement on metabolite quantification (Near et al 2020) and described by Gasparovic et al (2006). This method accounts for the differential water T1- and T2-relaxation in white matter, gray matter and CSF in the voxel to enable absolute metabolite measurements. The scripts for the correction were developed by (DeMayo et al., 2023), and are publicly available (https://github.com/HarrisBrainLab/TissueCorrections). However, this approach does not address differences in metabolite concentrations between white matter and gray matter (as suggested in (Gasparovic et al., 2018)) as this ratio is not agreed upon for all metabolites (and may change with development). For this reason, the gray matter tissue fraction was included as a covariate in analyses.

The gray matter tissue fraction of each MRS voxel was calculated as the fraction of voxel volume composed of gray matter divided by total tissue within the voxel (sum of the fractions of voxel volume for gray matter (GM) and white matter (WM; formula: fGM/[fGM + fWM]). The midline location of the ACC voxel was set to exclude white matter to the extent possible. To ensure minimal white matter in the ACC voxel, tissue fraction outliers were identified using boxplots and Rosner’s test using the *EnvStats R* package (Millard, 2013), and two datasets with tissue fraction <.88 were excluded from analysis (both from the unexposed group). Given that the LTP voxel included a substantial mix of gray and white matter in all participants, tissue fraction was applied as a covariate in analyses to account for individual differences in tissue composition of the voxels to ensure that group differences reflect differences in metabolites and not in tissue content of the voxels.

Code for processing and quantification of MRS is available here: https://github.com/developmental-neuroimaging-lab/mrs

### Cognitive assessments

Participants’ executive function and pre-reading abilities were measured with the NEuroPSYchological Assessment - Second Edition (NEPSY-II; Korkman et al., 2007), and the caregiver-report Behavior Rating Inventory of Executive Function (BRIEF; Gioia et al., 2000) or BRIEF-Preschool; (BRIEF-P; Gioia et al., 2003) using age-appropriate forms (BRIEF-P: ages 2-5 years, BRIEF: ages ≥6 years), most often administered on the same day or within two weeks of the MRI scan and always within six months. Cognitive data was available for a sub-set of participants in each cohort; EF skills (BRIEF/BRIEF-P and NEPSY-II Statue subtest) were examined in Cohort 1, and pre-reading skills (NEPSY-II Phonological Processing and Speeded Naming subtests) were examined in Cohort 2. Demographic characteristics of the sub-samples included in analysis of cognitive data are included in the Supplementary Information (Table S2). In addition, most participants completed the Wechsler Preschool and Primary Scale of Intelligence - Fourth Edition (WPPSI-IV; Wechsler, 2012) at their first study visit as a measure of global cognitive function. We report WPPSI-IV Full Scale IQ (FSIQ) standard scores in Tables 1 & 2 for sample characterization.

#### NEPSY-II

Inhibitory control (motor inhibition) was assessed via the Statue subtest of the NEPSY-II. This subtest is designed to test inhibitory control in young children, and it is the only NEPSY-II sub-test in the Attention and Executive Functioning domain that can be administered to children ages 3-4 years (and can be administered up to age 6 years). Participants were asked to stand still with their eyes closed and inhibit responses to sound distractors. A researcher observed the child while they tried to stand motionless with eyes closed for a 75 second period. A perfect score would indicate that the child remained still with eyes closed the entire time with increasingly lower scores indicating more errors (movement, eyes opened) during the task. Test-retest reliability (Pearson’s R coefficient) of the Statue Total Scaled Score is ∼.8 (Brooks et al., 2009).

Key pre-reading skills were assessed via the Phonological Processing (PP) and Speeded Naming (SN) subtests of the NEPSY-II. PP measured the child’s ability to segment and manipulate phonemes, and SN measured the child’s ability to rapidly identify pictured objects and colors.

Performance on NEPSY-II sub-tests is indexed by scaled scores based on population norms (mean=10, SD=3) for child age. Higher scores indicate better performance.

#### BRIEF and BRIEF-P

Global executive functioning was assessed using the BRIEF or BRIEF-P as appropriate for each child’s age (PAE: n=6 BRIEF, n=10 BRIEF-P; Unexposed: n=2 BRIEF, n=22 BRIEF-P). These measures are parent-report measures of naturalistic executive function. Children’s executive functioning abilities across various sub-domains are summarized in the Global Executive Composite (GEC) T-score, normed for child age. GEC T-scores are scaled based on population norms (mean=50, SD=10) for child age. Higher GEC T-scores indicate poorer performance.

### Data analysis

### Group differences in metabolite levels

Statistical analysis was performed in RStudio (RStudio Team, 2020) using *R* version 4.3.1 (R Core Team, 2016). Group differences in metabolite levels were tested using linear mixed effects models (*lme4* and *lmerTest* packages; Bates et al., 2015; Kuznetsova et al., 2017) with random intercept by participant to account for repeated measures within participants. Metabolite differences between exposed and unexposed children were tested with the formula:

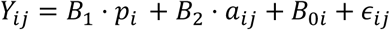

Where *Y_ij_* = jth metabolite measurement for the ith subject, *B*_1_ = coefficient for PAE status, *p_i_* = the ith subject’s PAE status (0 or 1), *B*_2_ = coefficient for age, *a_ij_* = ith subject’s age at time of jth scan, *B*_0*i*_= subject-specific y-intercept, and *∊*_*ij*_ = random error.

Age at MRI was included as a covariate in models in which a significant main effect of age on metabolite level was observed (tCho, tNAA, both cohorts); age was dropped from models in which its inclusion did not improve model fit relative to the more parsimonious model. We did not include sex in the models because our prior MRS research in this age range has not shown sex differences in metabolite concentrations. Post-hoc sensitivity analyses in sub-samples matched on number of scans, age, and sex, and in the full samples including sex as a covariate are reported in the Supplementary Information (Tables S3 & S4). Tissue fraction was included as a covariate for LTP analyses only. Random effects were dropped from models in which random effects variance = 0 (Glx, both cohorts). False discovery rate (FDR; Benjamini & Hochberg, 1995) correction was applied separately in each cohort to correct for multiple comparisons; we report both uncorrected and corrected results.

### Metabolite associations with EF and pre-reading skills

Associations between metabolites and cognitive scores were tested for metabolites that showed significant group differences, with separate models for each metabolite.

For Cohort 1 (ACC), associations between metabolites (tCho and Glx) and EF measures (age-normed BRIEF/BRIEF-P GEC T-scores and NEPSY-II Statue sub-test scaled scores) and PAE-by-metabolite interactions were tested using linear models with the formula:

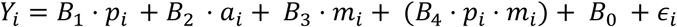

Where *Y_i_* = EF score for the ith subject, *B*_1_ = coefficient for PAE status, *p_i_* = the ith subject’s PAE status (0 or 1), *B*_2_ = coefficient for age, *a_i_ = ith subject’s age at cognitive assessment, *B*_3_ = the coefficient for metabolite measurement, *m_i_* = the ith subject’s metabolite measurement, *B*_4_ = the coefficient for the interaction between PAE status and metabolite measurement, *B*_0_ = intercept, and *∊*_*i*_* = random error.

Age at cognitive assessment was included as a fixed effect for the NEPSY-II Statue analysis, but was not included for the BRIEF/BREIF-P analysis because age was not associated with BRIEF/BRIEF-P GEC T-scores and its inclusion did not improve model fit.

For Cohort 2 (LTP), associations between tCho and age-normed pre-reading scores (PP or SN) and PAE-by-metabolite interactions were tested using linear mixed effects models with the formula:

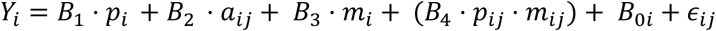

Where *Y_i_* = pre-reading score for the ith subject, *B*_1_ = coefficient for PAE status, *p_i_* = the ith subject’s PAE status (0 or 1), *B*_2_ = coefficient for age, *a_ij_* = ith subject’s age at time of jth visit, *B*_3_ = the coefficient for metabolite measurement, *m_ij_* = the ith subject’s metabolite measurement (adjusted for tissue fraction by residualization) at time of jth scan, *B*_4_ = the coefficient for the interaction between PAE status and metabolite measurement, *B*_0*i*_= subject-specific y-intercept, and *∊*_*ij*_ = random error.

Code for statistical analysis is available at: https://osf.io/4zkd8/?view_only=f7526d82205044feb07796410a937ca5

## Results

### Metabolite differences in PAE vs. unexposed children

Cohort 1: The PAE group had elevated levels of tCho (*beta* = .226, 95% CI [.04, .41], *p* = .016, *p_FDR_* = .045) and Glx (*beta* = 1.474, 95% CI [.19, 2.76], *p* = .025, *p_FDR_*= .054) in the ACC relative to unexposed children (Table 1, Figure 2), though the Glx effect is marginal. No significant group differences were found for tNAA, tCr or mI in the ACC. Measures of MRS data quality (Linewidth and SNR) differed significantly between groups, with children with PAE generally having lower quality data, but post-hoc analyses including linewidth and SNR as covariates in the models showed no significant effects of these metrics on tCho and Glx, and the pattern of elevated tCho and Glx in the PAE group was maintained in these models; see Supplementary Information for details (Table S5). Moreover, poorer data quality (greater linewidth and lower SNR) is expected to result in *lower* (rather than higher) estimates of metabolite concentrations, which is opposite to the observed pattern in our data, further supporting the robustness of these results.

**Figure 2.**
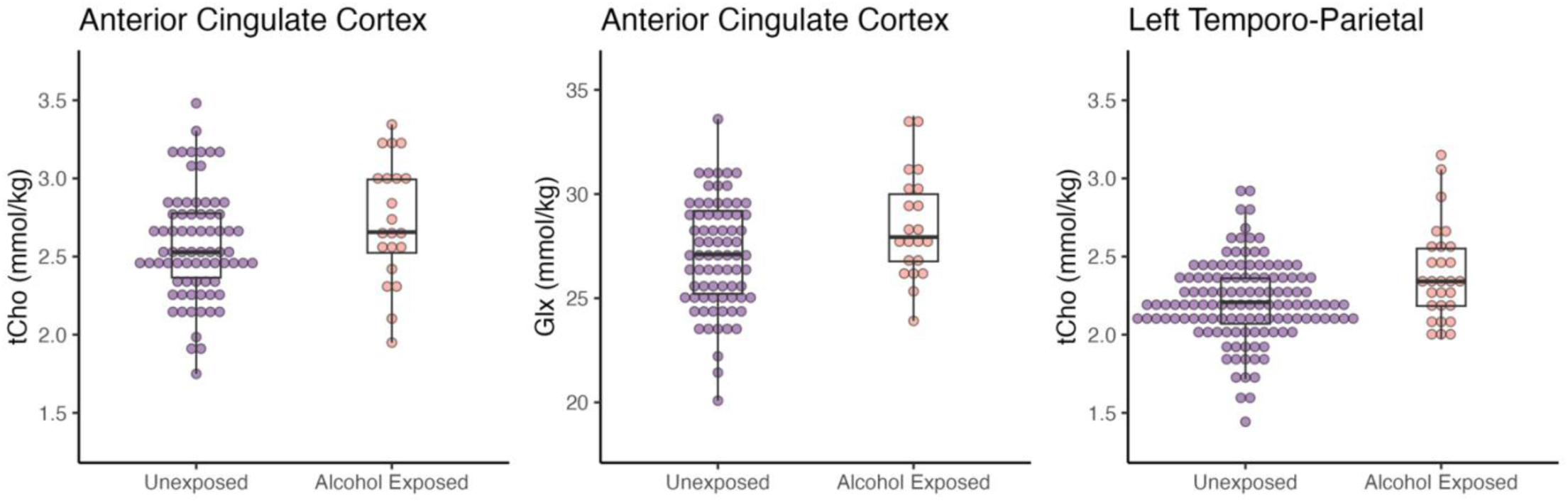
Metabolite differences by alcohol exposure group shown by box plots indicating group means and quartiles overlaid on individual data points. Group differences in metabolite levels in the ACC (Cohort 1; left and middle panels) and LTP (Cohort 2; right panel) shown for metabolites with significant effects before FDR correction; only the tCho effect in the ACC remained significant after FDR correction for multiple comparisons (*p*=.016, *p_FDR_*=.045). Individual data points plotted (purple=unexposed, orange=PAE).

**Table 1.** Descriptive statistics and group differences in metabolites, quality metrics, and cognitive skills for Cohort 1. Unstandardized beta coefficients from linear mixed effects models reported unless otherwise noted. N observations reported, with the exception of Family Income which is reported for N participants. Significant results are indicated in bold (uncorrected level) and * (FDR corrected). Family Income=Total Household Income Range reported by caregiver at time of participation; Maternal Education=years of postsecondary education reported by mother in the home at time of participation. N participants and median income range reported in table, group difference tested using Wilcoxon Rank Sum tests for family income and two-sample t-tests for maternal education and WPPSI-IV FSIQ.

| Cohort 1: ACC | Unexposed Children |  |  | Prenatal Alcohol Exposure |  |  | Group Difference |  |  |
| --- | --- | --- | --- | --- | --- | --- | --- | --- | --- |
| Full sample | N obs. | Mean | SD | N obs. | Mean | SD | $\beta$ | $p$ | $p_{FDR}$ |
| Female/Male | 36/40 | - | - | 11/10 | - | - | $\chi^2=.026$ | .873 | - |
| Family Income | 64 | \$150,000 - 174,999 | | 16 | \$75,000 - 99,999 | | **$W=912.5$** | **<.001** | - |
| Maternal Education | 68 | 5.65 | 3.14 | 15 | 2.87 | 1.81 | **$t=4.615$** | **<.001** | - |
| WPPSI-IV FSIQ | 48 | 108.5 | 10.88 | 20 | 92.35 | 13.4 | **$t=4.773$** | **<.001** |  |
| Age at MRI (years) | 76 | 4.51 | 0.95 | 21 | 5.34 | 1.12 | **0.889** | **<.001** | **.002\*** |
| tNAA (mmol/kg) | 76 | 15.84 | 1.16 | 21 | 15.89 | 1.1 | -0.0179 | 0.543 | .710 |
| tCr (mmol/kg) | 76 | 12.8 | 1.1 | 21 | 12.83 | 1.16 | 0.095 | 0.757 | .804 |
| tCho (mmol/kg) | 75 | 2.58 | 0.35 | 21 | 2.73 | 0.38 | **0.226** | **0.016** | **.045\*** |
| Glx (mmol/kg) | 76 | 27.02 | 2.57 | 21 | 28.46 | 2.52 | **1.474** | **0.025** | **.054** |
| ml (mmol/kg) | 74 | 7.57 | 1.56 | 19 | 7.55 | 1.95 | -.010 | 0.982 | .982 |
| Tissue Fraction (GM/[GM+WM]) | 76 | 0.96 | 0.02 | 21 | 0.97 | 0.03 | 0.009 | 0.126 | .214 |
| Linewidth (NAA) | 76 | 7.39 | 3.04 | 21 | 10.25 | 3.01 | **2.766** | **<.001** | **.002\*** |
| SNR (NAA) | 76 | 56.67 | 9.62 | 21 | 46.72 | 7.44 | **-9.616** | **<.001** | **.001\*** |
| Cognitive Sub-sample | N obs. | Mean | SD | N obs. | Mean | SD | $\beta$ | $p$ | $p_{FDR}$ |
| Age at Assessment (years) | 25 | 4.37 | 0.76 | 17 | 5.54 | 1.15 | **1.168** | **<.001** | **.002\*** |
| NEPSY-II Statue Total Scaled Score | 25 | 10.56 | 2.72 | 17 | 9.76 | 4.09 | .479 | 0.691 | .792 |
| BRIEF/BRIEF-P GEC T-score | 22 | 49.23 | 11.56 | 16 | 73.44 | 17.29 | **23.544** | **<.001** | **.001\*** |

Cohort 2: The PAE group had elevated levels of tCho (*beta* = .135, 95% CI [.02, .25], *p* = .02, *p_FDR_* = .10) in the LTP relative to the unexposed group (Table 2, Figure 2), though this effect did not pass FDR correction for multiple comparisons. No significant group differences were found for tNAA, tCr, Glx or mI in the LTP.

**Table 2.**
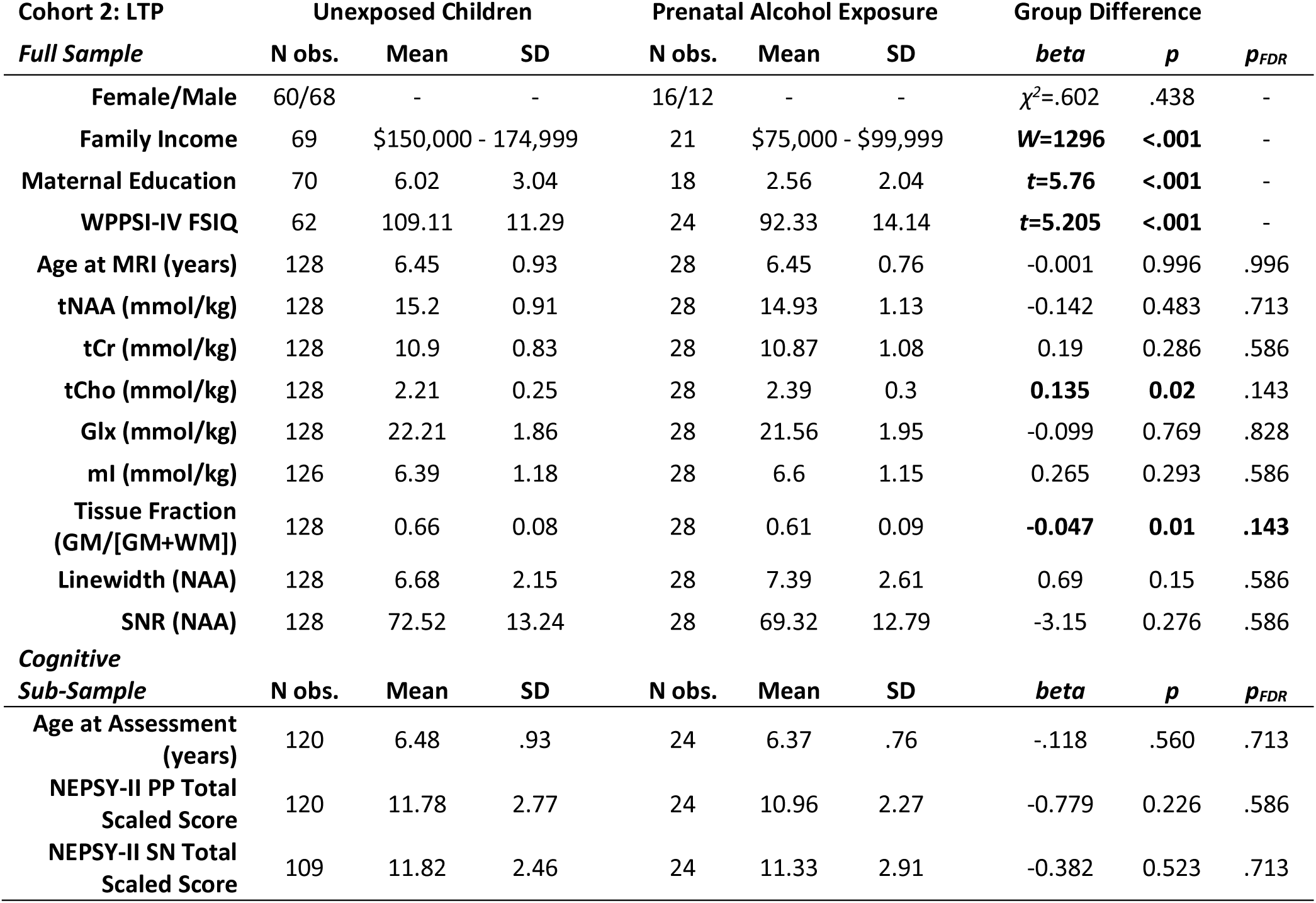
Descriptive statistics and group differences in metabolites, quality metrics, and cognitive skills for Cohort 2. Unstandardized beta coefficients from linear mixed effects models reported unless otherwise noted. N observations reported, with the exception of Family Income which is reported for N participants. Significant results are indicated in bold (uncorrected level). Family Income=Total Household Income Range reported by caregiver at time of participation; Maternal Education=years of postsecondary education reported by mother in the home at time of participation. N participants and median income range reported in table, group difference tested using Wilcoxon Rank Sum tests for family income and two-sample t-tests for maternal education and WPPSI-IV FSIQ.

## Executive functioning and pre-reading skill associations with PAE and metabolites

### Cohort 1 EF and Metabolites in the ACC

A regression model examining the effects of PAE status and Glx on inhibitory control measured via the NEPSY-II Statue Total Scaled Score revealed a significant interaction between PAE status and Glx (*beta* = - .83, 95% CI[-1.55, -.11], *p*=.026, *p_FDR_*=.054). Post-hoc analysis indicated a negative association between Glx and inhibitory control in children with PAE, but not unexposed children (Figure 3A). No significant associations between inhibitory control and tCho were found (*p*>.3).

**Figure 3.**
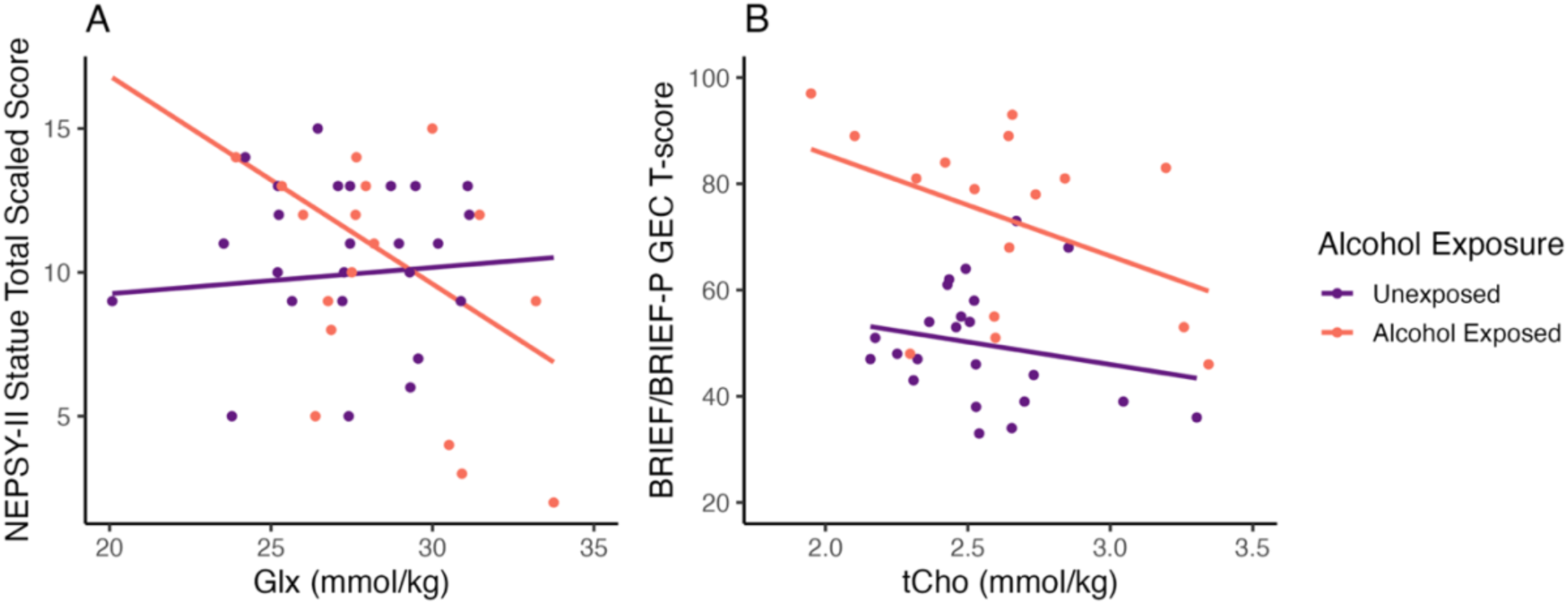
**Associations between EF measures and metabolites in ACC (Cohort 1).** A) Inhibitory control was associated with Glx in the ACC (Cohort 1, n=42) in the PAE group (orange) but not the unexposed group (purple). Inhibitory control indexed by NEPSY-II Statue Total Scaled Score. B) Associations between PAE status, ACC tCho levels, and EF (Cohort 1, n=39). Fit lines represent negative association between tCho and EF within each group for unexposed (red) and PAE (yellow). EF indexed by BRIEF/BRIEF-P GEC T-score: higher scores indicate more difficulties in executive functioning.

A regression model examining the effects of PAE status and tCho on EF indexed by the BRIEF-II/BRIEF-P GEC T-score indicated that children with PAE had worse executive functioning relative to unexposed children (*beta* = 24.92, 95%CI[16.08, 33.76], *p*<.001, *p_FDR_*<.001), and lower tCho was associated with worse executive functioning in both groups (*beta*=-15.07, 95%CI[-29.12, -1.01], *p*=.036, *p_FDR_*=.066) (Table 1; Figure 3B). No significant group-by-metabolite interactions were found, and BRIEF/BRIEF-P GEC scores were not significantly associated with Glx (*p*>.3).

### Cohort 2 Pre-reading skills and metabolites in the LTP

Pre-reading skills (PP and SN) did not show any significant associations with tCho in the LTP, or alcohol exposure status. Groups performed similarly on PP and SN measures and group means were in the average to above-average range (Table 2).

## Discussion

In this study, we report the first evidence of PAE-related brain metabolite differences in early childhood, showing elevated tCho and Glx in children with PAE relative to unexposed children. Group differences in tCho in the ACC were significant after FDR correction for multiple comparisons, while group differences in Glx in the ACC and tCho in the LTP were significant at the uncorrected level. These findings offer insight to neurochemical disruptions that correspond to differences in brain structure and function observed in PAE across childhood. Further, we show associations between brain metabolites and measures of executive function, one of the most commonly observed areas of cognitive/behavioral difficulty in individuals with PAE.

### Metabolite differences in PAE vs. unexposed children

We found higher tCho in both the ACC and LTP of children with PAE compared to unexposed children, though the effect in the LTP (cohort 2) did not pass correction for multiple comparisons. tCho levels are sensitive to the presence of all types of cell membranes and are associated with myelin content and fiber density (Perdue et al., *under review*). Thus, our findings likely indicate widespread alterations in tissue microstructure and phospholipid metabolism across brain regions and tissue types (gray and white matter) in children with PAE. The ACC voxel examined in Cohort 1 included >90% gray matter, so our results likely reflect more complex cortical architecture in the children with PAE, possibly as a result of reduced/delayed pruning of synapses and more dense packing of neurites. In Cohort 2, the LTP voxel contained a mix of gray and white matter, and elevated tCho was present, even after accounting for individual differences in tissue composition of the MRS voxels. Together, the tCho effects in our study may reflect differences in both gray and white matter microstructure, including neurite density and myelin content.

Notably, our finding of *higher* tCho in PAE groups contrasts with prior evidence that showed *lower* tCho in older children and adolescents with PAE. This may point to altered developmental trajectories of phospholipid metabolism, myelination, and cortical development in PAE, consistent with findings from structural brain imaging studies (Gimbel et al., 2023; Kar et al., 2022). In fact, no differences in tCho were found in an MRS study of neonates with PAE (Howells et al., 2016), suggesting that alterations in tCho may emerge later in postnatal development and change over childhood. We recently showed that tCho levels decrease in early childhood until about 9 years of age in unexposed children (Perdue et al., 2023), so elevated levels of tCho in early childhood in our cohorts with PAE likely reflect a developmental delay in the tCho decline that accompanies myelination and cortical refinement. Our present results show larger group differences in tCho levels in Cohort 1 (children ages 3-6 years old) compared to Cohort 2 (children ages 5-8 years old). Although we cannot fully compare the two cohorts due to differences in brain regions examined, this suggests that elevated tCho observed in our study may be driven by younger children, and a reversal of this effect may occur in middle-late childhood when tCho levels tend to stabilize. This fits with studies in older children, where *reduced* tCho is observed in groups with PAE (Astley et al., 2009; du Plessis et al., 2014; Gonçalves et al., 2009). Our lab previously showed a similar developmental pattern in white matter microstructure, where differences in early childhood showed an unexpected pattern opposite to prior research in older samples. However, longitudinal data showed that these early differences diminished or reversed direction with age around middle childhood, indicating altered developmental trajectories in children with PAE that help explain discrepant findings between younger and older children (Kar et al., 2021, 2022). However, it should be noted that methodological differences (location of MRS voxel placement, quantification methods [water-referencing vs. creatine ratios], and methods of correcting for tissue content) could also contribute to mixed findings across MRS studies. Longitudinal research tracking tCho across childhood in children with PAE is needed to help reconcile findings in younger vs. older populations.

Glx was elevated in the ACC (Cohort 1), but not the LTP (Cohort 2). Given that glutamate is the primary excitatory neurotransmitter of the central nervous system and primary contributor to the Glx signal in MRS, over-abundance of Glx in the ACC is consistent with hypothesized hyperexcitability in fronto-striatal circuits in PAE (Bariselli & Lovinger, 2021). This hypothesis is supported by findings of altered functional activity in frontal brain regions, including the ACC, which are thought to be associated with EF difficulties in individuals with PAE (Fryer et al., 2007; Kodali et al., 2017; Ware et al., 2015). Concerning mechanism, higher measurements of Glx by MRS likely reflect an excess of excitatory synapses, indicating a delay or failure of synaptic pruning in the ACC and potential for hyperexcitability. Prior results for Glx in individuals with PAE have been somewhat mixed, with higher Glx in the cerebellum associated with PAE in older children (du Plessis et al., 2014), but lower Glx in parietal white matter in newborns with PAE (Howells et al., 2016). Prior studies of Glx have also shown mixed findings in developmental trajectories, and it is likely that Glx changes non-linearly with age, showing increases in infancy/early childhood followed by decreases from late childhood/adolescence into early adulthood (Blüml et al., 2013; Ghisleni et al., 2015; Holmes et al., 2017; Perdue et al., 2023; Perica et al., 2022; Thomson et al., 2024; Volk et al., 2019). This could result in elevated or reduced Glx in groups with PAE at different developmental stages. Alternatively, mixed findings across studies could arise from methodological differences in metabolite quantification, brain regions targeted, and methods of tissue correction. Nonetheless, longitudinal research in this area will be crucial to uncover how PAE-related differences in brain metabolites unfold across development.

### Associations between metabolites and executive functioning

Our results showing that increased Glx is associated with EF difficulties in children with PAE support the role of hyperexcitability in altered behavior and cognition in the context of PAE. In a subset of participants who completed the NEPSY-II Statue task, higher Glx (i.e., higher excitability) in the ACC was associated with poorer motor inhibitory control. This effect was specific to the PAE group, which indicates a particular role of Glx and hyperexcitability and inefficient inhibitory control in children with PAE. Alterations in Glx may reflect dysfunction in the cycling of glutamate, glutamine, and the inhibitory neurotransmitter, GABA, which is needed to maintain the excitatory-inhibitory balance for efficient functioning (Bak et al., 2006; Sears & Hewett, 2021). This is consistent with preclinical research that indicates that PAE may disrupt the corticalstriatal excitatory-inhibitory balance by increasing glutamatergic activity *via* altered synapse formation and enhanced AMPA receptor function, and reducing GABA-ergic function *via* disrupted migration of inhibitory neurons (Bariselli & Lovinger, 2021). The absence of an association between Glx and inhibitory control in the unexposed group may indicate that unexposed children have more optimal Glx levels and intact glutamate-glutamine-GABA cycling. Thus, it may be that intact inhibitory systems and glutamate-glutamine-GABA cycling in unexposed individuals with relatively high Glx (similar levels to the PAE group) are better able to maintain the excitatory-inhibitory balance, resulting in no net effect of Glx on inhibitory control task performance. However, as the inhibitory neurotransmitter, GABA, was not measured in this study, nor in any prior human study of PAE, future research is needed to explicitly compare inhibitory systems in groups with and without PAE.

In addition to the association between Glx and inhibitory control, we observed a main effect of tCho on global EF indexed by the BRIEF/BRIEF-P GEC, such that higher tCho in the ACC was associated with better global EF. This finding was paired with a strong main effect of PAE status, such that children with PAE displayed more difficulties with EF relative to unexposed children. tCho levels in gray matter likely reflect packing density of cell bodies and neurites as well as ongoing phospholipid metabolism to maintain healthy tissue (Rae, 2014). Thus, higher tCho in the ACC may indicate that there are more neurons and synaptic connections available for top-down cognitive control, and that this architecture confers a benefit for EF within groups with and without PAE, despite overall EF and tCho differences between groups. This interpretation is supported by evidence that linked cortical morphometry of the ACC (greater surface area) to faster reaction time on a cognitive control task in typically developing children (Fjell et al., 2012), and another study that linked poorer inhibition to reduced ACC surface area in adolescents with PAE (Migliorini et al., 2015).

Together, the observed associations between metabolite levels in the ACC and EF suggest that PAE-related alterations in excitability, metabolism, and cortical microstructure are related to cognitive and behavioral difficulties in children with PAE. In contrast to EF, prereading skills (phonological processing or speeded naming) were spared in our sample (Cohort 2) and not associated with metabolite levels in the LTP. This is not surprising, as EF is known to be particularly impaired in PAE and findings related to language and reading skills in PAE have been mixed (Subramoney et al., 2018). Language and literacy impairments may depend on the amount and timing of alcohol exposure (O’Leary et al., 2013), and difficulties in this domain become apparent later in the acquisition of reading skills, so further research is needed to fully characterize PAE-related associations between brain metabolites and cognition.

### Limitations

Our study has several limitations that should be considered. First, due to the challenges of recruiting and acquiring data from children with PAE, we were not always able to obtain a full, detailed birth history, nor specific amounts and timing of PAE (though PAE was confirmed in all cases). Nonetheless, co-occurring conditions such as pre-term birth or low birth weight could explain some of the findings. Participants with and without PAE were not matched on sociodemographic characteristics such as socioeconomic status and home environment, which may contribute to observed differences in brain and cognition. Further research in samples with and without PAE matched on birth-related and socio-demographic factors are needed to attempt to isolate the effects of alcohol exposure on brain metabolites and cognition. In addition, although most children were awake during scanning, we were unable to control for whether children were awake or asleep in our analysis because this was not tracked in detail at the individual level. Future research should consider the sleep/wake status of participants to account for possible sleep-related fluctuations in metabolites. Importantly, no sedation was used in this study. Finally, we did not examine sex or gender differences in our study, but such effects may emerge over development and future research should consider possible sex and gender differences associated with brain chemistry in groups with and without PAE.

## Conclusion

Alcohol disrupts multiple aspects of brain development and functioning. Here we show persistent effects of PAE on brain metabolites in early childhood, with implications for cognition and behavior. This evidence can support the development of interventions to modulate cortical hyperexcitability and improve EF in children with PAE. Future research combining multimodal neuroimaging techniques with MRS in PAE is needed to provide further insight to the metabolic mechanisms underlying altered brain structure and function in PAE.

## Statements and Declarations

### Author Contributions

**Meaghan V. Perdue:** conceptualization, methodology, formal analysis, writing – original draft, writing – review & editing, data visualization; **Mohammad Ghasoub:** data curation, formal analysis, writing – original draft, writing – review & editing; **Madison Long:** data curation, writing – original draft, writing – review & editing; **Marilena M. DeMayo:** methodology, writing – review & editing; **Tiffany K. Bell:** methodology, writing – review & editing; **Carly A. McMorris:** conceptualization, writing – review & editing; **Deborah Dewey:** conceptualization, writing – review & editing, funding acquisition; **W. Ben Gibbard:** writing – review & editing; **Christina Tortorelli:** writing – review & editing; **Ashley D. Harris:** methodology, writing – review & editing, supervision; **Catherine Lebel:** conceptualization, methodology, writing – review & editing, funding acquisition, supervision.

### Data Availability

Code used for pre-processing and quantification of MRS data is available at https://github.com/developmental-neuroimaging-lab/mrs; Data processing and analysis code are available at https://osf.io/4zkd8/?view_only=f7526d82205044feb07796410a937ca5; Data is available upon request to the corresponding author.

## Supplementary Information

**Table S1.**
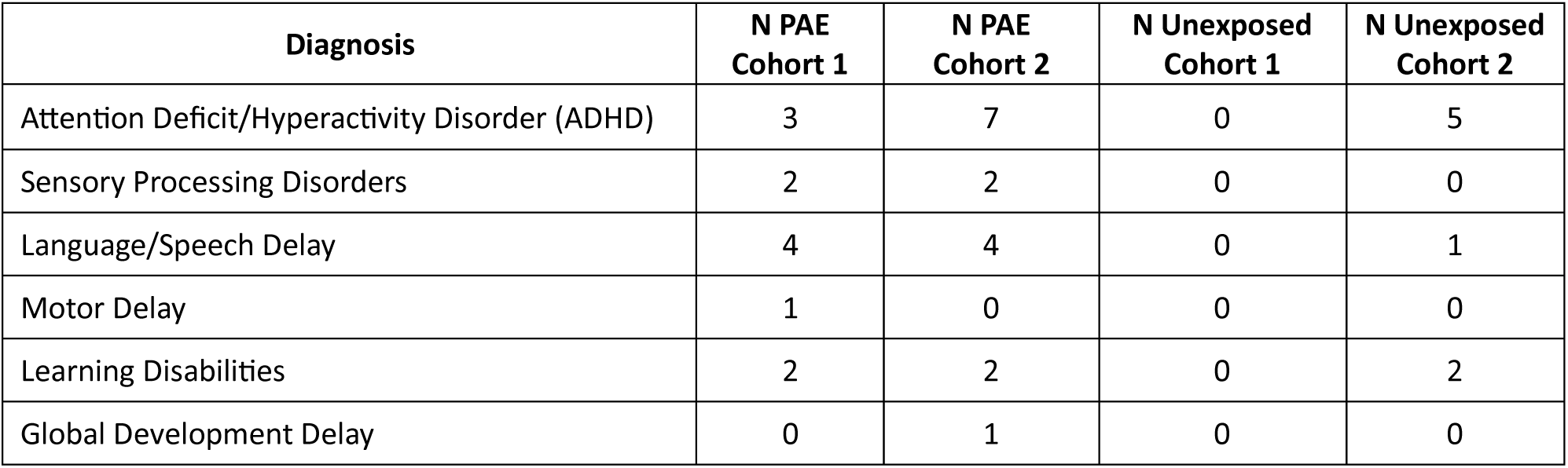
Neurodevelopmental conditions diagnosed in PAE participants in each cohort, based on parent report.

**Table S2.**
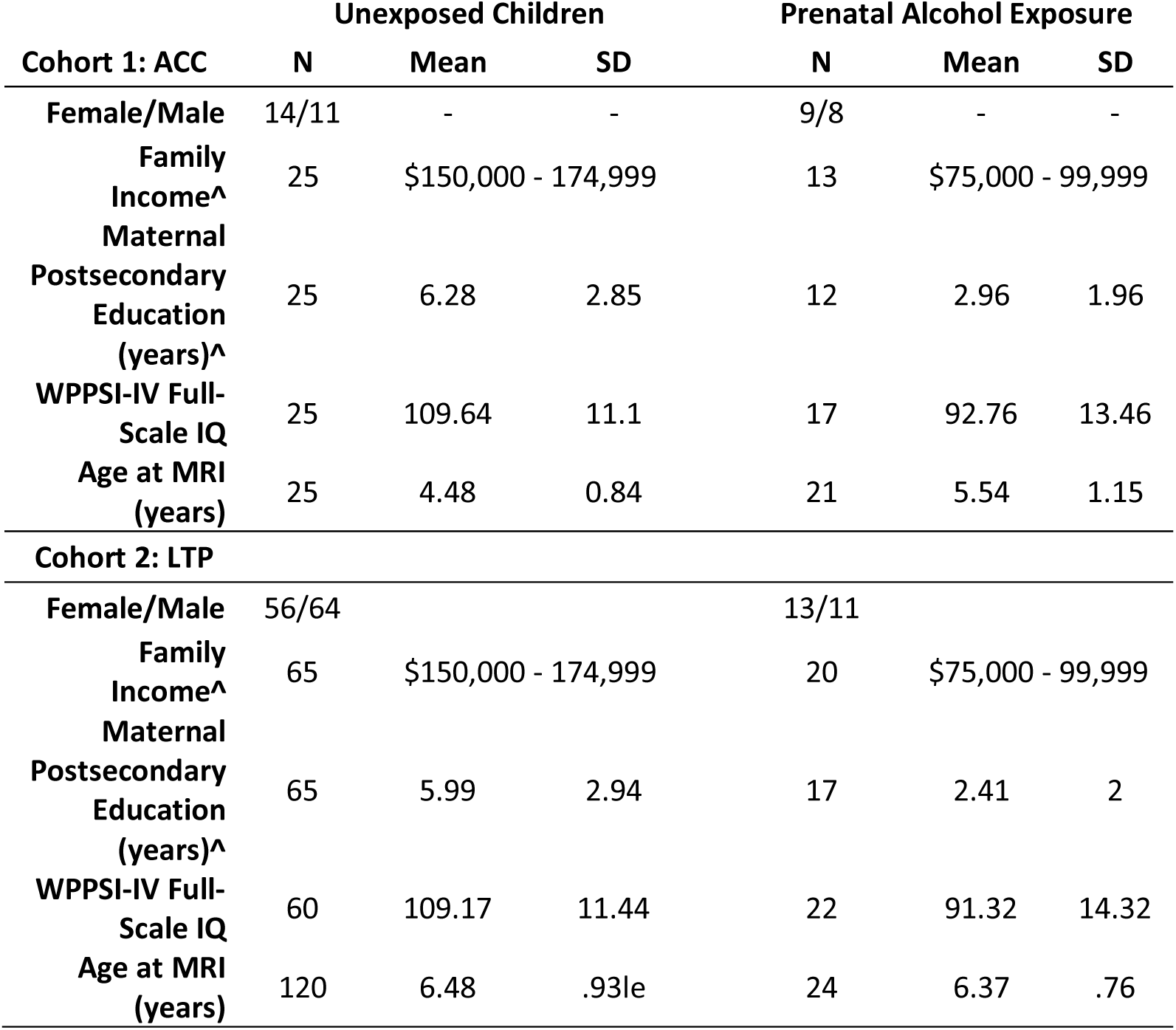
Demographic characteristics of sub-samples included in analysis of cognitive abilities. N individuals reported for Cohort 1 (no repeated individuals) and N individuals reported for Cohort 2 Family Income, Maternal Education and WPPSI-IV FSIQ variables; N observations reported for Cohort 2 sex and age at MRI.

**Table S3.**
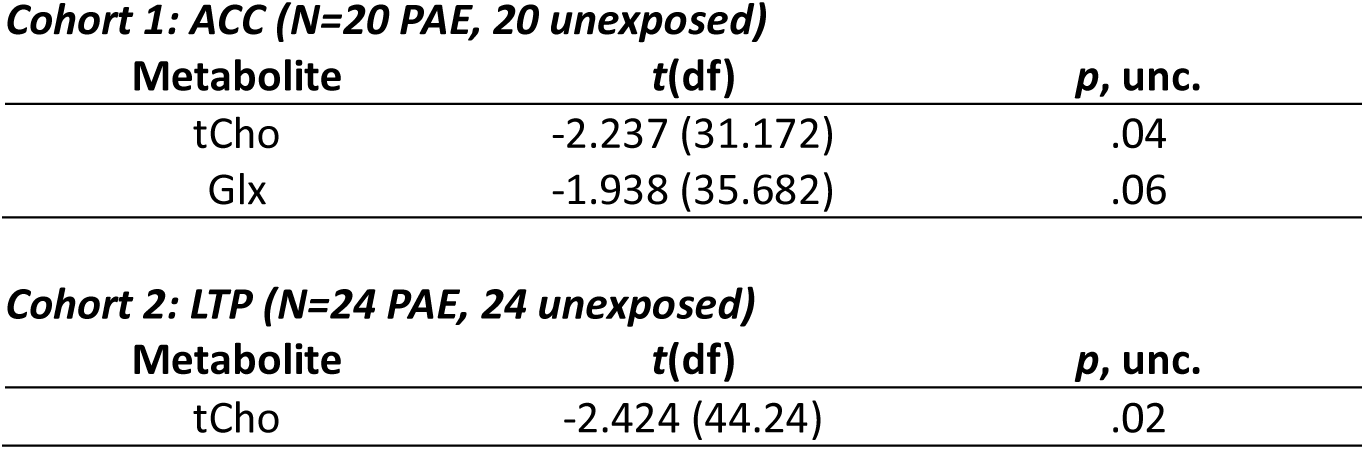
Post-hoc analysis of metabolite differences in groups matched on number of scans (no repeated scans within subjects), age, sex, and tissue fraction. The first available scans in the ACC and LTP for PAE individuals were used as the base datasets for matching. Results of two-sample T-tests in the matched samples are consistent with our findings from the main linear mixed effects models on the full samples.

**Table S4.**
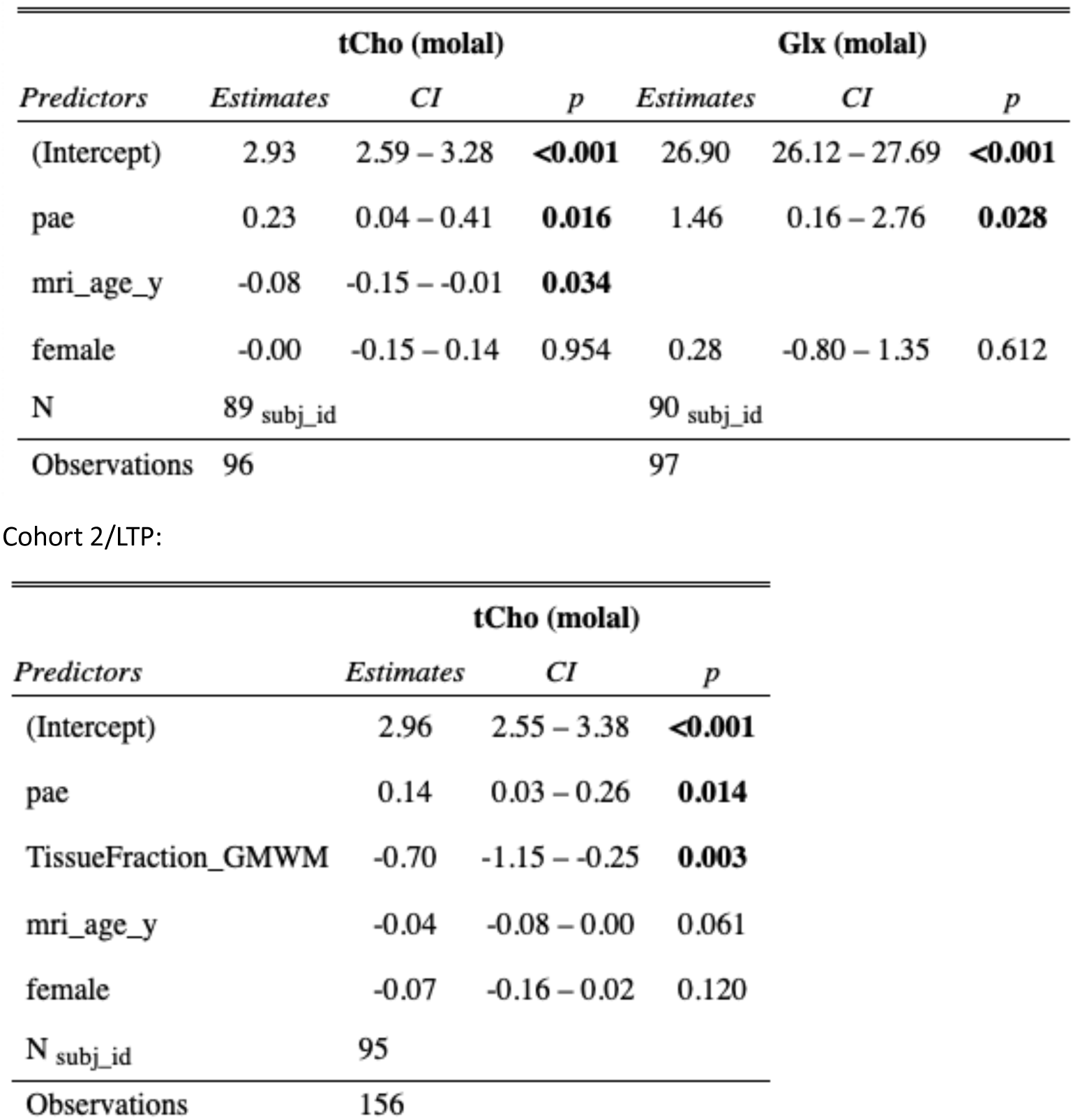
Model summaries for post-hoc sensitivity analyses for LME models including sex as a covariate. Cohort 1/ACC:

**Table S5.**
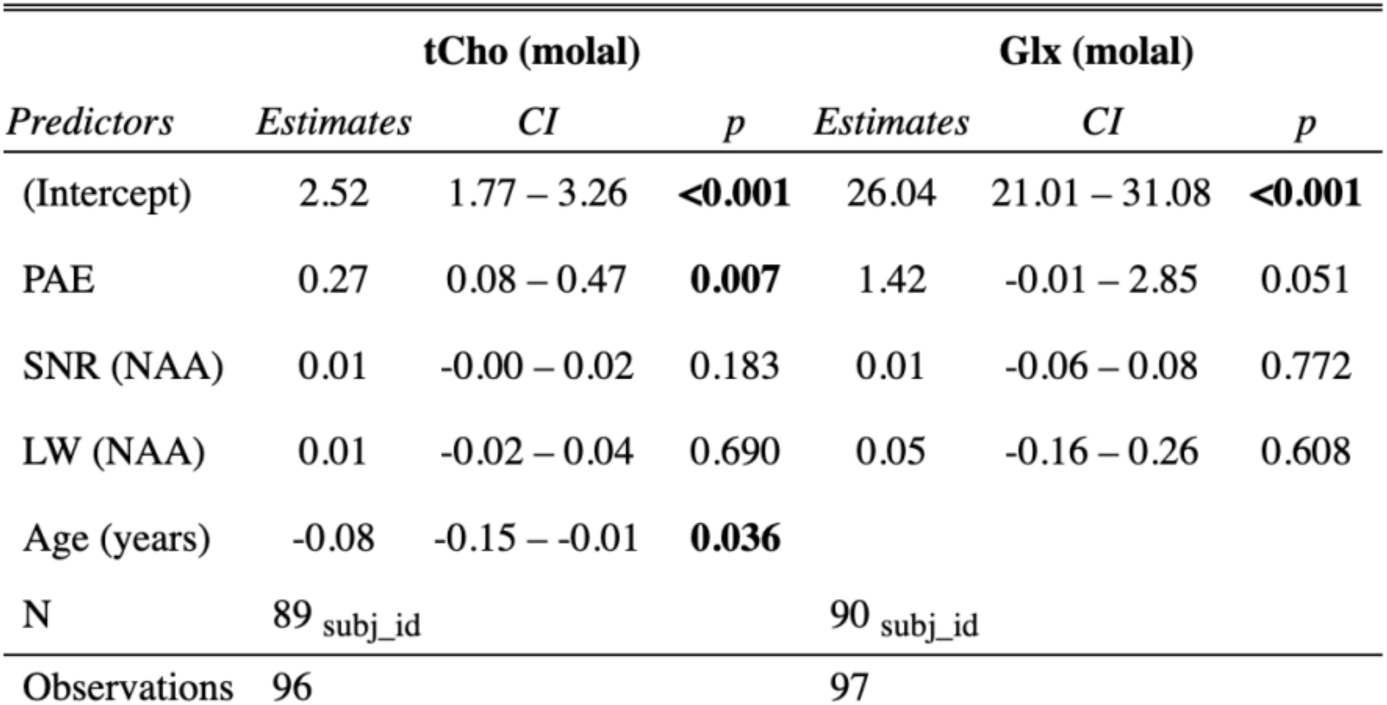
Model summaries for post-hoc sensitivity analysis of Cohort 1 (ACC) including MRS quality metrics (SNR and linewidth [LW] of the NAA peak) as covariates.

