## Supplementary Information for "Altered markers of brain metabolism and excitability are associated with executive functioning in young children exposed to alcohol *in utero*"

**Table S1.** Neurodevelopmental conditions diagnosed in PAE participants in each cohort, based on parent report.

| Diagnosis | N PAE Cohort 1 | N PAE Cohort 2 | N Unexposed Cohort 1 | N Unexposed Cohort 2 |
|---|---|---|---|---|
| Attention Deficit/Hyperactivity Disorder (ADHD) | 3 | 7 | 0 | 5 |
| Sensory Processing Disorders | 2 | 2 | 0 | 0 |
| Language/Speech Delay | 4 | 4 | 0 | 1 |
| Motor Delay | 1 | 0 | 0 | 0 |
| Learning Disabilities | 2 | 2 | 0 | 2 |
| Global Development Delay | 0 | 1 | 0 | 0 |

**Table S2.** Demographic characteristics of sub-samples included in analysis of cognitive abilities. N individuals reported for Cohort 1 (no repeated individuals) and N individuals reported for Cohort 2 Family Income, Maternal Education and WPPSI-IV FSIQ variables; N observations reported for Cohort 2 sex and age at MRI.

| | Unexposed Children | | | Prenatal Alcohol Exposure | | |
|---|---|---|---|---|---|---|
| **Cohort 1: ACC** | **N** | **Mean** | **SD** | **N** | **Mean** | **SD** |
| **Female/Male** | 14/11 | - | - | 9/8 | - | - |
| **Family Income^** | 25 | $150,000 - 174,999 | | 13 | $75,000 - 99,999 | |
| **Maternal Postsecondary Education (years)^** | 25 | 6.28 | 2.85 | 12 | 2.96 | 1.96 |
| **WPPSI-IV Full-Scale IQ** | 25 | 109.64 | 11.1 | 17 | 92.76 | 13.46 |
| **Age at MRI (years)** | 25 | 4.48 | 0.84 | 21 | 5.54 | 1.15 |
| **Cohort 2: LTP** | | | | | | |
| **Female/Male** | 56/64 | | | 13/11 | | |
| **Family Income^** | 65 | $150,000 - 174,999 | | 20 | $75,000 - 99,999 | |
| **Maternal Postsecondary Education (years)^** | 65 | 5.99 | 2.94 | 17 | 2.41 | 2 |
| **WPPSI-IV Full-Scale IQ** | 60 | 109.17 | 11.44 | 22 | 91.32 | 14.32 |
| **Age at MRI (years)** | 120 | 6.48 | .93le | 24 | 6.37 | .76 |

**Table S3.** Post-hoc analysis of metabolite differences in groups matched on number of scans (no repeated scans within subjects), age, sex, and tissue fraction. The first available scans in the ACC and LTP for PAE individuals were used as the base datasets for matching. Results of two-sample T-tests in the matched samples are consistent with our findings from the main linear mixed effects models on the full samples.

***Cohort 1: ACC (N=20 PAE, 20 unexposed)***

| Metabolite | *t*(df) | *p*, unc. |
|---|---|---|
| tCho | -2.237 (31.172) | .04 |
| Glx | -1.938 (35.682) | .06 |

***Cohort 2: LTP (N=24 PAE, 24 unexposed)***

| Metabolite | *t*(df) | *p*, unc. |
|---|---|---|
| tCho | -2.424 (44.24) | .02 |

**Table S4.** Model summaries for post-hoc sensitivity analyses for LME models including sex as a covariate.

Cohort 1/ACC:

| | tCho (molal) | | | Glx (molal) | | |
|---|---|---|---|---|---|---|
| *Predictors* | *Estimates* | *CI* | *p* | *Estimates* | *CI* | *p* |
| (Intercept) | 2.93 | 2.59 – 3.28 | **<0.001** | 26.90 | 26.12 – 27.69 | **<0.001** |
| pae | 0.23 | 0.04 – 0.41 | **0.016** | 1.46 | 0.16 – 2.76 | **0.028** |
| mri_age_y | -0.08 | -0.15 – -0.01 | **0.034** | | | |
| female | -0.00 | -0.15 – 0.14 | 0.954 | 0.28 | -0.80 – 1.35 | 0.612 |
| N | 89 $_{subj\_id}$ | | | 90 $_{subj\_id}$ | | |
| Observations | 96 | | | 97 | | |

Cohort 2/LTP:

| | tCho (molal) | | |
|---|---|---|---|
| *Predictors* | *Estimates* | *CI* | *p* |
| (Intercept) | 2.96 | 2.55 – 3.38 | **<0.001** |
| pae | 0.14 | 0.03 – 0.26 | **0.014** |
| TissueFraction_GMWM | -0.70 | -1.15 – -0.25 | **0.003** |
| mri_age_y | -0.04 | -0.08 – 0.00 | 0.061 |
| female | -0.07 | -0.16 – 0.02 | 0.120 |
| N $_{subj\_id}$ | 95 | | |
| Observations | 156 | | |

**Table S5.** Model summaries for post-hoc sensitivity analysis of Cohort 1 (ACC) including MRS quality metrics (SNR and linewidth [LW] of the NAA peak) as covariates.

| | **tCho (molal)** | | | **Glx (molal)** | | |
|---|---|---|---|---|---|---|
| *Predictors* | *Estimates* | *CI* | *p* | *Estimates* | *CI* | *p* |
| (Intercept) | 2.52 | 1.77 – 3.26 | **<0.001** | 26.04 | 21.01 – 31.08 | **<0.001** |
| PAE | 0.27 | 0.08 – 0.47 | **0.007** | 1.42 | -0.01 – 2.85 | 0.051 |
| SNR (NAA) | 0.01 | -0.00 – 0.02 | 0.183 | 0.01 | -0.06 – 0.08 | 0.772 |
| LW (NAA) | 0.01 | -0.02 – 0.04 | 0.690 | 0.05 | -0.16 – 0.26 | 0.608 |
| Age (years) | -0.08 | -0.15 – -0.01 | **0.036** | | | |
| N | $89_{\text{subj\_id}}$ | | | $90_{\text{subj\_id}}$ | | |
| Observations | 96 | | | 97 | | |
